# Analysis of SARS-CoV-2 Recombinant Lineages XBC and XBC.1 in the Philippines and Evidence for Delta-Omicron Co-infection as a Potential Origin

**DOI:** 10.1101/2023.04.12.534029

**Authors:** Elcid Aaron R. Pangilinan, John Michael C. Egana, Renato Jacinto Q. Mantaring, Alyssa Joyce E. Telles, Francis A. Tablizo, Carlo M. Lapid, Maria Sofia L. Yangzon, Joshua Jose S. Endozo, Karol Sophia Agape R. Padilla, Jarvin E. Nipales, Lindsay Clare D.L. Carandang, Zipporah Mariebelle R. Enriquez, Tricia Anne U. Barot, Romano A. Manlimos, Kelly Nicole P. Mangonon, Ma. Exanil L. Plantig, Shiela Mae M. Araiza, Jo-Hannah S. Llames, Kris P. Punayan, Rachelle P. Serrano, Anne M. Drueco, Honeylett T. Lagnas, Philip A. Bistayan, Aristio C. Aguilar, Joie G. Charisse Apo, Yvonne Valerie D. Austria, Niña Francesca M. Bustamante, Alyssa Jamila R. Caelian, Rudy E. Fernandez, Xerxanne A. Galilea, Marielle M. Gamboa, Clarence Jane A. Gervacio, Zyrel V. Mollejon, Joshua Paul N. Pineda, Kristel B. Rico, Jan Michael C. Yap, Ma. Celeste S. Abad, Benedict A. Maralit, Marc Edsel C. Ayes, Eva Maria Cutiongco-de la Paz, Cynthia P. Saloma

## Abstract

We report the sequencing and analysis of 60 XBC and 114 XBC.1 SARS-CoV-2 lineages detected in the Philippines from August to September 2022, which are regarded as recombinant lineages of the BA.2 Omicron and B.1.617.2 Delta (21I Clade) variants. The sequences described here place the Philippines as the country with the earliest and highest number of XBC and XBC.1 cases within the included period. Majority of the detected cases were sampled from the adjacent Davao and Soccskargen regions in southern Philippines, but have also been observed at lower proportions in other regions of the country. Time-scaled phylogenetic analysis with global samples from GISAID reaffirms the supposed root of XBC-like cases from the Philippines. Furthermore, the apparent clustering of some foreign cases separate from those collected in the country suggests several occurrences of cross-border transmissions resulting in the spread of XBC-like lineages within and among those countries. The consensus mutation profile shows regions harboring mutations specific to either the Omicron BA.2 or Delta B.1.617.2 lineages, supporting the recombinant nature of XBC. Finally, alternative allele fraction pattern and intrahost mutation analysis revealed that a relatively early case of XBC collected in March 2022 is likely to be an active co-infection event. This suggests that co-infection of Omicron and Delta was already occurring in the Philippines early in 2022, facilitating the generation of recombinants that may have further evolved and gained additional mutations enabling its spread across certain local populations at a later time.

**Author summary:** More recently, various lineages of the SARS-CoV-2 virus, the causative agent COVID-19 pandemic, have been observed to form recombinant lineages, further expanding the ways by which the virus can evolve and adapt to human interventions. Therefore, a large part of biosurveillance efforts is dedicated to detecting and observing new lineages, including recombinants, for early and effective control. In this paper, we present an analysis of 174 XBC and XBC.1 cases detected in the Philippines between August and September of 2022 which contextualize these cases as some of the earliest reported cases of this hybrid lineage. We show that when compared to cases from other countries collected at a similar time, the earliest cases of the XBC lineage are from the Philippines. Additionally, when samples were reclassified following an update of Pangolin, a tool for assigning SARS-CoV-2 lineages to samples, we found two samples of interest reclassified as XBC pointing to a potential origin via co-infection events occurring as early as March of 2022.

## Introduction

Recombination is known to occur in RNA viruses such as HIV-1 [1], SARS-CoV [2], and now, SARS-CoV-2 [3]. In March 2021, a provision in the naming convention for confirmed SARS-CoV-2 recombinant lineages within the PANGO dynamic nomenclature system [4] states all top-level lineages that are recombinants have a prefix that begins with X [5].

As of December 2022, there are around 75 notable recombinant lineages listed in the pango-designation lineage notes [6], including XBC and XBC.1 first described in late September 2022. XBC is believed to be a recombinant lineage from the BA.2 Omicron and the B.1.617.2 (21I Clade) Delta parentage, previously suspected as a Delta saltation because of highly divergent sequences particularly in the spike region resembling that of BA.2 sublineage. The presence of a high number of mutations was later hypothesized to be a by-product of recombination [7]. On the other hand, XBC.1 was designated to have an additional S:L452M mutation not present in the original XBC lineage [8]. According to the sequences submitted in the EpiCoV database of the Global Initiative for Sharing All Influenza Data (GISAID) [9], the Philippines is the country with the earliest occurrence and the highest number of detected XBC and XBC.1 cases from September 2022 to present.

In the Philippines, two major COVID-19 surges were associated with the Omicron variant of concern. The first surge started during the start of the year 2022 with a steep increase happening in mid-January peaking in late January followed by a steady decline and plateauing in March (Fig 1A). This surge resulted in the highest number of recorded daily case counts in the country and was primarily driven by the BA.2.3 Omicron lineage. The second surge was mainly linked to the BA.5.2 Omicron sublineages. In contrast with the first surge, this second surge exhibited a slower increase in cases starting in mid-June and peaking at the start of August. Afterwards, the number of confirmed cases decreased marginally but did not completely subside, stabilizing at around 2,000 daily confirmed cases beginning in September. During this plateau, cases in Davao (Region 11) did not decrease substantially after peaking at 125 in mid-August. The cases instead immediately stabilized at 100 daily new cases (Fig 1B). Similarly, cases in Soccsksargen (Region 12) continued to slowly increase even as the number of daily confirmed cases nationwide appeared to plateau (Fig 1C).

**Fig 1.**
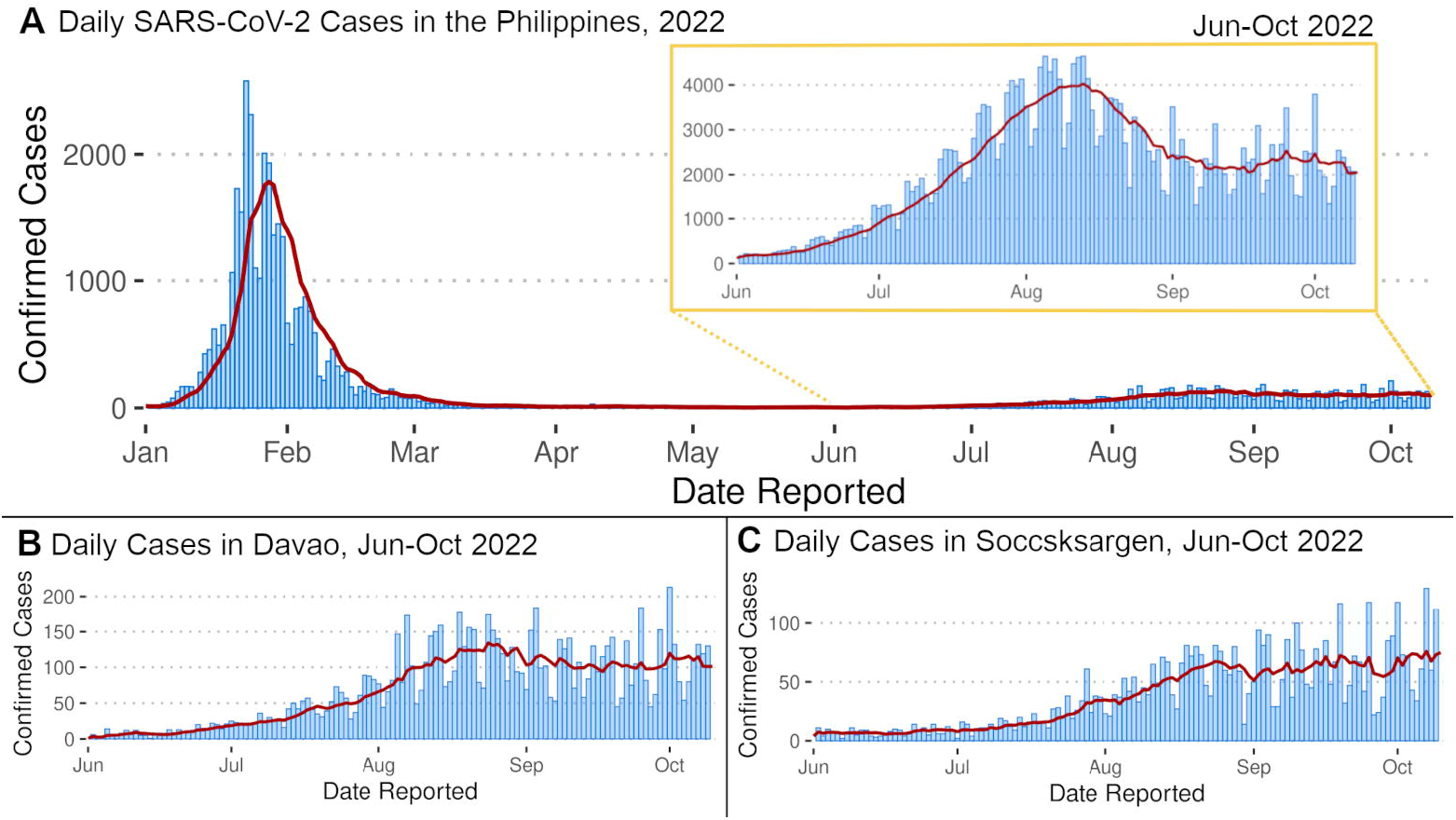
Daily new reported SARS-CoV-2 cases, smoothed with a 7-day moving average. Daily new cases in (A) the Philippines and in the southern regions of (B) Davao and (C) Soccsksargen, using data obtained from the Department of Health. More than three months after the first Omicron surge at the beginning of 2022, the Philippines observed another increase in reported SARS-CoV-2 cases (inset) that started to decline in August but continued to plateau throughout September. During the plateau, a small increasing trend can be observed in Davao and Soccsksargen.

The plateau in daily case counts observed in the aforementioned regions coincided with detected samples from our genomic biosurveillance data being assigned to the recombinant XBC lineage and its corresponding sublineages, particularly XBC.1. This led us to further investigate these samples in terms of their spatio-temporal distribution and genomic profiles.

## Results and Discussion

### Local Distribution of XBC and XBC.1 Cases (Aug - Sep 2022)

We report the sequencing and analyses of 60 XBC and 114 XBC.1 Philippine

SARS-CoV-2 cases collected throughout August to September 2022, most of which were from southern Mindanao (Fig 2). Mindanao is a major island in the Philippine archipelago consisting of six regions: Zamboanga Peninsula (Region 9), Northern Mindanao (Region 10), Davao (Region 11), Soccsksargen (Region 12), Caraga (Region 13) and Bangsamoro Autonomous Region of Muslim Mindanao (BARMM). We initially observed an increasing proportion of cases classified as the ancestral B.1.617.2 Delta variant in these regions, which were later re-assigned to the then-emerging variant XBC and its sublineage XBC.1 (Fig 3). These lineages are described by pango-lineages [7] as recombinant lineages of the BA.2 Omicron and B.1.617.2 Delta (21I Clade) variants.

**Fig 2.**
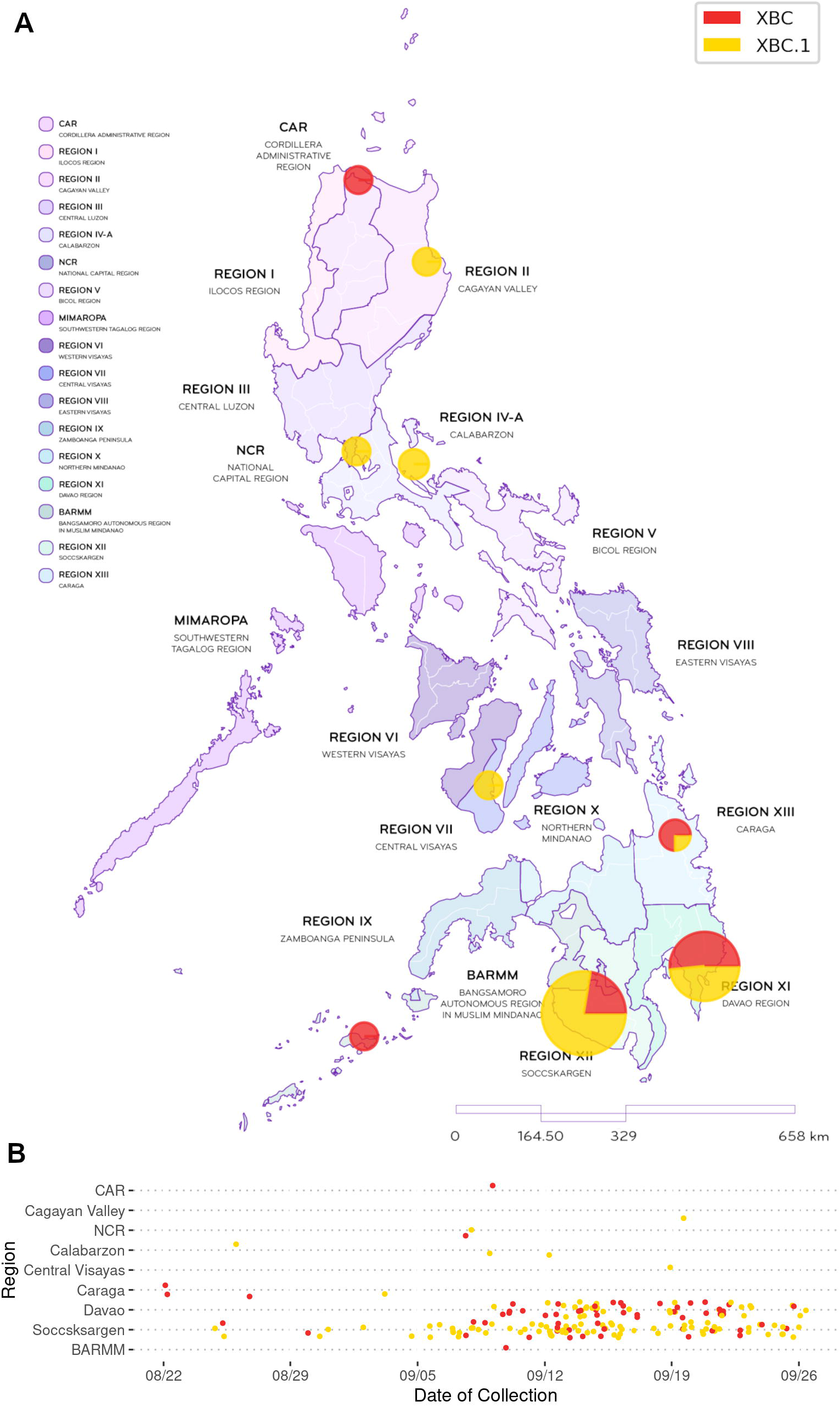
Map and jitter plot of the distribution of detected XBC and XBC.1 cases across the Philippines from Aug-Sep 2022. Initial cases of XBC in August 2022 were observed mainly in southern Philippine regions (Caraga, Soccsksargen), followed by an increase in detected XBC.1 cases in Davao and Soccsksargen. The two lineages were observed to have circulated primarily in the adjacent regions of Davao and Soccsksargen in September 2022. A few cases of XBC.1 were also detected in other regions suggesting that it has spread across the country.

**Fig 3.**
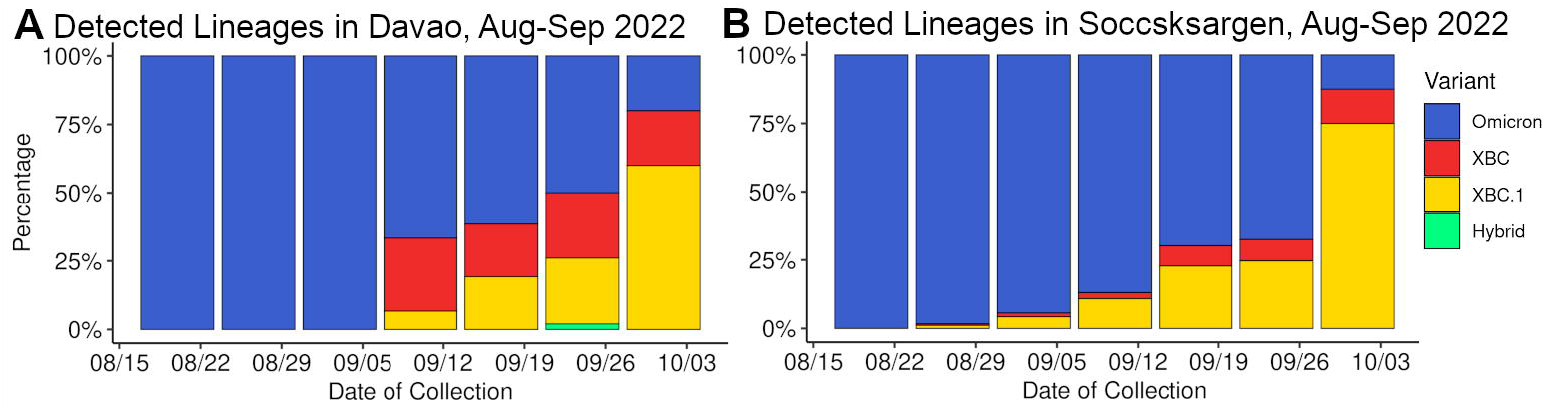
Stacked percentage bar chart showing the weekly proportion of detected lineages in Mindanao regions from Aug-Sep 2022. (A) Davao weekly proportions and (B) Soccsksargen weekly proportions, where most of the XBC and XBC.1 cases were detected. As recorded COVID-19 cases nationwide plateaued around September 2022, XBC and XBC.1 were observed to be increasing in proportion particularly in the regions of Davao and Soccsksargen.

Although mainly observed in the southern regions of Philippines, local cases of XBC and its sublineage XBC.1 have already been detected in other parts of the country (Fig 2; Supplementary Table S1). The high proportion of these cases in Davao and Soccsksargen may be explained by the high land traffic volumes in these adjacent regions [10], while the further spread in other parts of the country may be due to Davao having the busiest airport in the island of Mindanao, carrying around 1.6 million passengers domestically from January to August 2022 [11].

### Phylogenetic Analysis of XBC and XBC.1 Cases (Aug - Sep 2022)

Phylogenetic analysis of XBC and XBC.1 cases sequenced in the country with foreign cases obtained from GISAID when colored by lineage (Fig 4A) shows that although occasional mixing was observed in XBC cases clustering within the XBC.1 clade, the sublineage designation is generally well supported by the genomic tree. It is also worth noting here that cases originating from the Philippines (Fig 4B) predate the foreign cases, with the bulk of the earlier cases coming from Davao (XBC) and Soccsksargen (XBC.1) (Fig 4C) which may support the idea that the XBC-like lineages emerged in the Philippines before spreading to other regions of the globe. Nonetheless, the apparent clustering of some of the foreign cases separate from those from Philippines may also be indicative of several instances of cross-border transmissions resulting in the spread and local evolution of XBC-like lineages within and among those countries.

**Fig 4.**
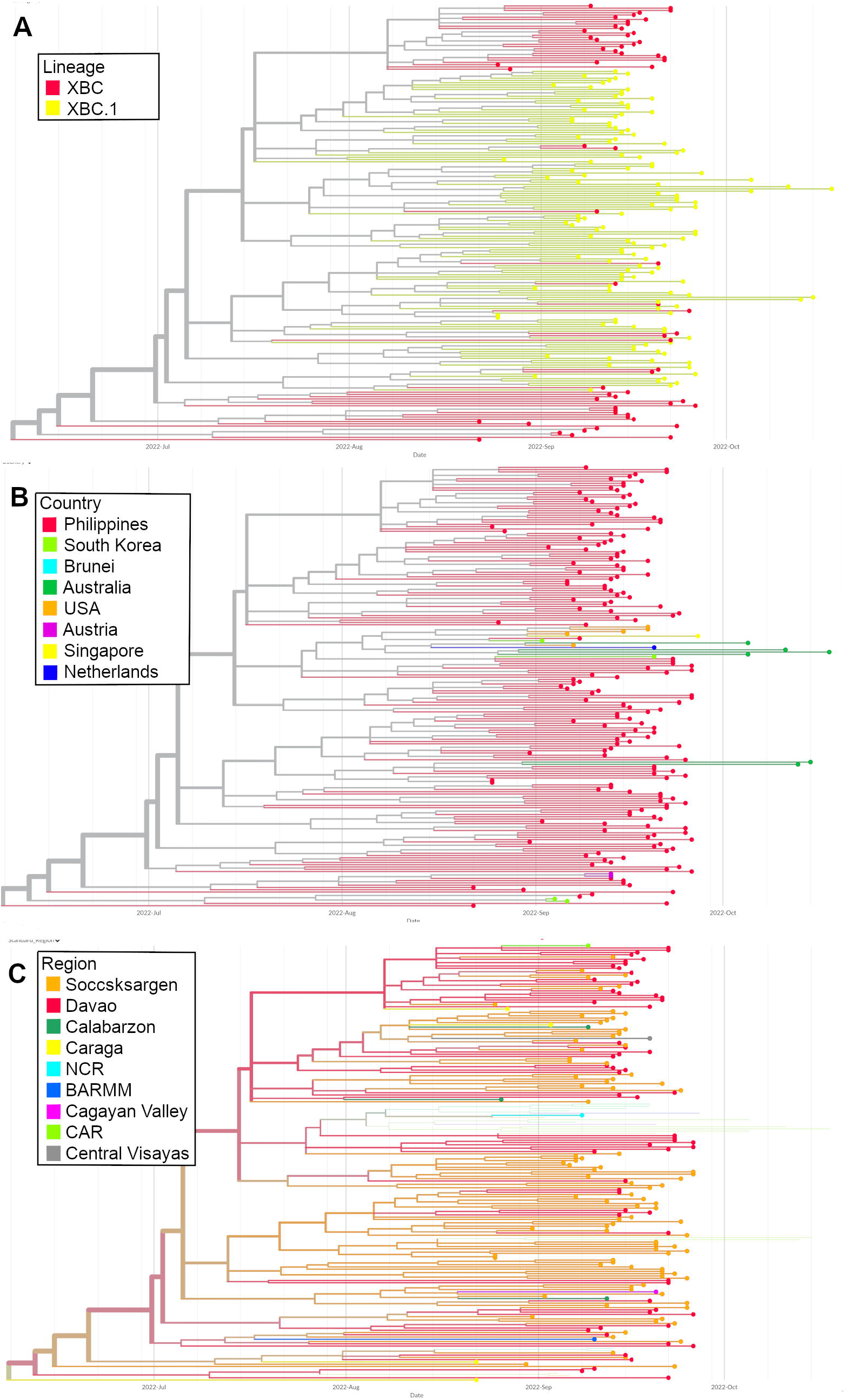
Phylogenetic analysis of 174 sequences sampled in the Philippines from August to September 2022. Time-scaled phylogenetic tree of locally sequenced XBC and XBC.1 cases supplemented with foreign sequences taken from GISAID colored by (A) lineage classification, (B) country of origin, and (C) region of origin in the Philippines. The tree colored by lineage shows occasional mixing of XBC-classified samples within the XBC.1 clade, but generally supports the sublineage designation. Based on country of origin, some of the sampled foreign XBC sequences formed clusters separate to those from the Philippines suggesting several instances of cross-border transmissions resulting in local spread within and among those countries. Looking at the region of origin in the Philippines, the XBC clusters are mainly samples from Davao whereas XBC.1 clusters are primarily from Soccsksargen – two adjacent regions in the southern part of the Philippines where the majority of XBC-like cases during the included period have been observed.

### Mutation Profile of Locally Detected XBC and XBC.1 Cases

Consistent with pango-lineage’s description of the lineages, local XBC and XBC.1 cases were found to harbor sections highly similar to those of the BA.2 Omicron and B.1.617.2 (21I clade) Delta variants (Fig 5). This would suggest XBC and XBC.1 as an Omicron-Delta recombinant with three breakpoints: between genomic positions 2790 (with ORF1ab:T842I) to 4184 (missing ORF1ab:G1307S), 22227 (with S:A222V) to 22674 (with S:S371F and missing S:L452R), and between 25469 (with S:N969K and missing ORF3a:S26L) to 26060 (missing ORF3a:T223I). This would imply that though Omicron greatly outcompeted Delta throughout the country towards the end of 2021, Delta-like lineages have continued to circulate in the country albeit at substantially lower proportions. Thus, there remained the possibility for co-infection and recombination between the co-circulating Delta and Omicron lineages.

**Fig 5.**
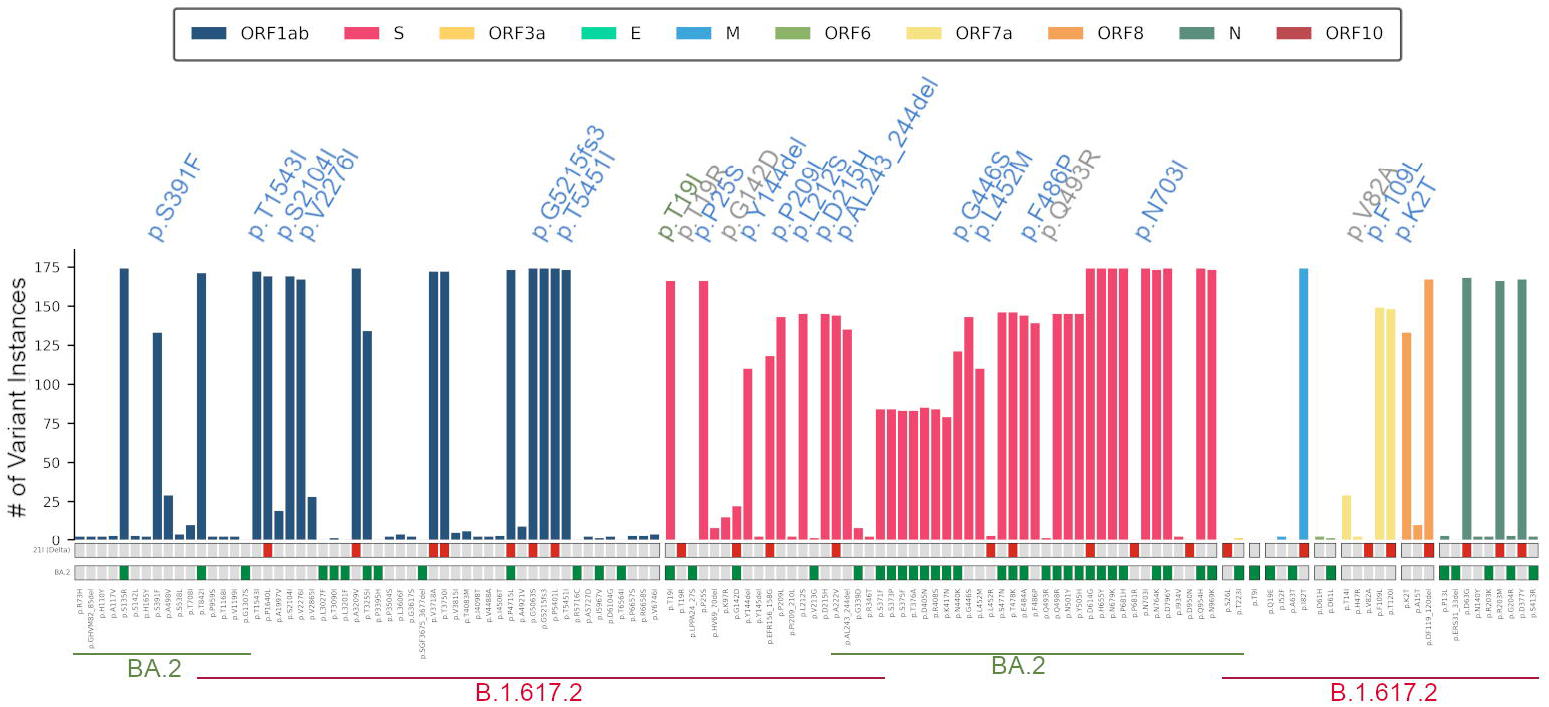
Consensus mutation profile of 174 local XBC and XBC.1 cases compared to the key mutations of BA.2 Omicron (green) and B.1.617.2 21I Delta Clade (red). These indicate the presence of mutations consistent with distinct sections of BA.2 and B.1.617.2. Highlighted are 18 distinct amino acid replacements and deletions (blue), and absences of key mutations (gray) compared to the BA.2 and B.1.617.2 sections. Additionally, though flanked by B.1.617.2 key mutations, the T19I Omicron mutation is found instead of the T19R Delta mutation. The separate consensus mutation profiles for the 60 XBC and the 114 XBC.1 are in Supplementary Fig S1.

Though a recombination is likely, an alternative explanation to the emergence of XBC is of convergent evolution. With regions of high similarity to BA.2 and B.1.617.2, these local XBC also harbor 18 amino acid replacements and deletions not frequently observed in BA.2 nor B.1.617.2 samples globally, including a frameshift insertion (ORF1ab:G5215fs3) (Fig 5). Four signature mutations were noticeably absent in the consensus mutation profile, while the S:T19I Omicron mutation is found instead of the S:T19R Delta mutation in a region supposedly from a Delta donor. Additionally, a section of mutations from the spike protein gene – from genomic position 22674 (S:S371F) to 22813 (S:K417N) – were detected at lower base call frequencies, which may also be due to primer amplification biases during enrichment.

### Evidence of a Pre-XBC Delta-Omicron Co-infection and Recombination in the Philippines

Interestingly, we have also detected two SARS-CoV-2 cases that have been reassigned as XBC (from a previous Delta classification) that were collected in March 2022, much earlier than most of the locally detected XBC-like cases. These samples, PH-PGC-113124 (collected 2022 March 20) and PH-PGC-113286 (collected 2022 March 31) were found to show recombination signals using 3SEQ v1.7 (p = 2E-12 and 1.1E-11, respectively) and Sc2rf with two primary breakpoints each: between 22200-22205 and 23525-23598, and 21618-21623 and 23604-23853, respectively (Fig 6A). This is in contrast to the consensus XBC sequence that has three breakpoint regions as described in the mutation profile analysis. We also note that for PH-PGC-113124, the recombinant region derived from the BA.2 Omicron sequence corresponding to the S gene is shorter than that of PH-PGC-113286, which is in turn shorter than that of the consensus XBC recombinant region. Because of the notable difference in the breakpoint regions, we hypothesize that these two samples represent earlier recombination events that eventually gave rise to the XBC lineage.

**Fig 6.**
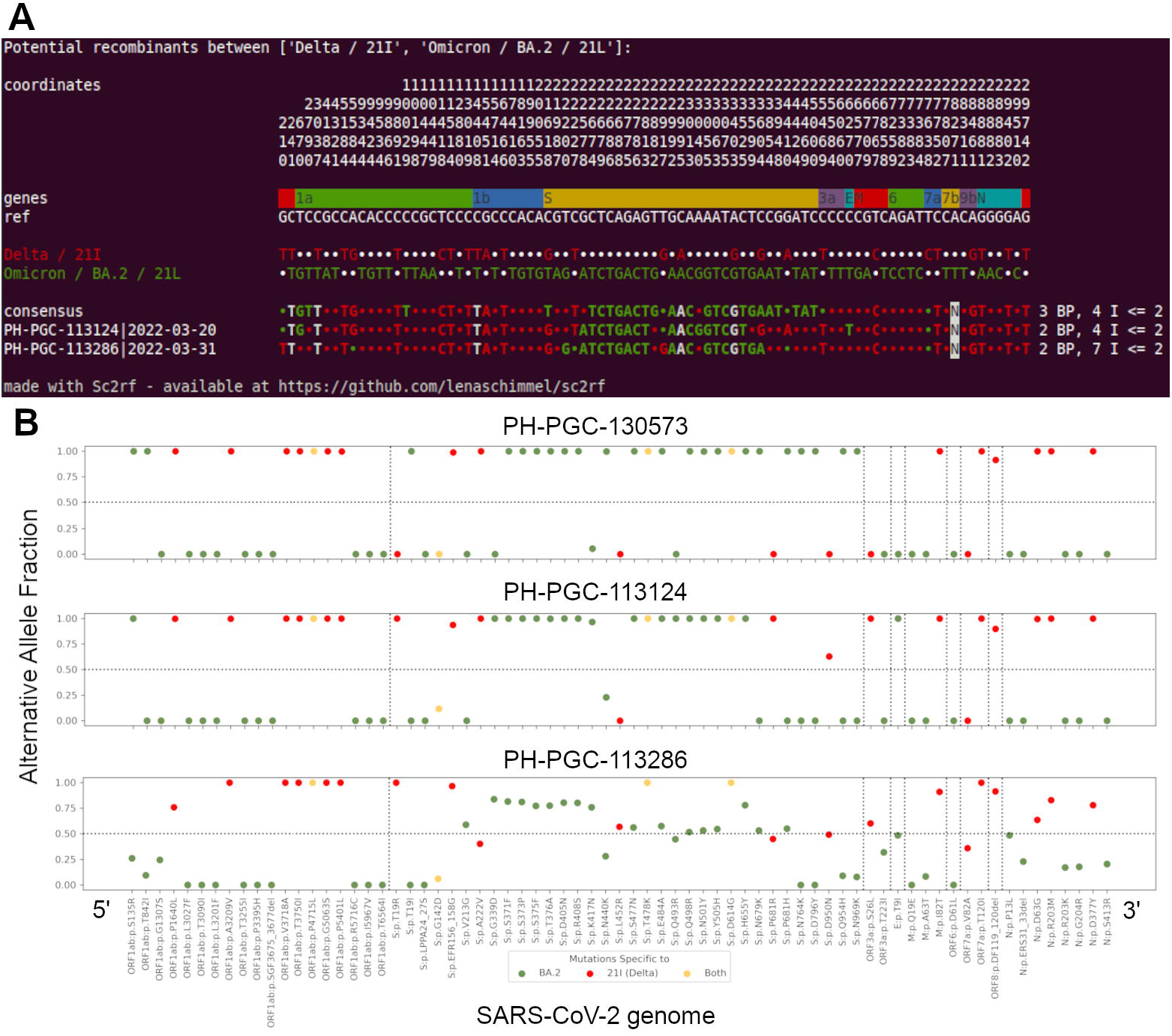
Recombination analysis of the two previously classified Delta samples collected in March 2022, PH-PGC-113124 and PH-PGC-113286, which have been reassigned as XBC recombinants. (A) Breakpoint analysis of the two samples along with the consensus sequence of the XBC variant reveals a slight discrepancy of the breakpoints in the spike region of the genome. Omicron signature mutations in the spike region of PH-PGC-113124 starts at position 22,578 up to 23,525; while PH-PGC-113286 has the signature Omicron mutations starting from position 22,220 up to 23,604. It is also worth noting that only two breakpoints were detected for the two samples compared to three for the consensus XBC sequence. (B) Alternative allele fraction analysis of a representative consensus XBC sequence (top), PH-PGC-113124 (middle), and PH-PGC-113286 (bottom). The mutations plotted are that of key mutations for BA.2 Omicron (green), B.1.617.2 21I Delta Clade (red), or mutations shared between them (yellow). These mutations were plotted in order of position in the SARS-CoV-2 genome, with dashed vertical lines separating the different gene regions. Mutations on the representative consensus sequence and PH-PGC-113124 are shown to either have nearly 1.0 or 0.0 alternative allele fractions, showing distinct sections of BA.2 and B.1.617.2. This follows the patterns of clonal recombinant sequences as described by Bolze et al. (2022). In contrast, PH-PGC-113286 is shown to initially have mutations with high allele fraction for Delta, which then substantially drops as the BA.2 defining mutations were detected. This is highly similar to a within-host recombination pattern during an active co-infection [3].

A closer investigation of these two early XBC cases revealed that the number of intrahost mutations (all mutations detected in a sample including those that are non-consensus) is much higher for PH-PGC-113286, harboring a total of 193 intrahost mutations, compared to that of PH-PGC-113124 with only 86. Note that the mean number of intrahost mutations for our XBC and XBC.1 samples are 78.2 and 99.7, respectively. This high number of intrahost mutations observed in PH-PGC-113286 could be explained by the presence of a co-infection, whereas PH-PGC-113124 is most likely already a case of clonal recombinant infection. Sequencing contamination may also be an explanation for the higher number of intrahost mutations, but we argue that this is highly unlikely as its parent lineage Delta B.1.617.2 had not been widely detected in our genomic surveillance during this period.

To further test the co-infection hypothesis, we performed an alternative allele fraction analysis as described in Bolze et al. [3] for the two early XBC samples (PH-PGC-113124 and PH-PGC-113286), as well as a representative from the more recently detected XBC cases already with three breakpoint regions, PH-PGC-130573 (Fig 6B). The PH-PGC-113124 sample is shown to have both the BA.2 and B.1.617.2 key mutations at nearly 100% allele fractions. This suggests that all the reads in the viral population of this sample are composed of a single recombinant variant [3]. The same pattern can be observed for the more recent XBC sample, indicating that both of these infections were already from a recombinant form of the virus to begin with.

In contrast, the PH-PGC-113286 sample contains BA.2 and B.1.617.2 key mutations supported by only a fraction of the reads. This indicates a mixture of full genome copies of multiple variants in the viral population, similar to the pattern indicative of co-infection [3]. In addition, PH-PGC-113286 begins with nearly 100% alternative allele fraction for Delta key mutations near the 5’ end, then drops and fluctuates to around 80% and 50% near the beginning of the spike protein gene – just as the mutations transition to signatures of the BA.2 lineage. This pattern is highly similar to that for a within-host recombination [3], suggesting that PH-PGC-113286 already harbors recombinants in some of the host cells while having an active co-infection.

These observations can serve as evidence of the occurrence of multiple Delta-Omicron co-infection cases in the Philippines prior to the emergence of the XBC recombinant lineage, providing insights to its possible origins. Nonetheless, we note that the products of these early recombination events likely evolved further and acquired additional mutations that enabled the XBC lineage to spread locally and better compete with other existing SARS-CoV-2 lineages.

## Conclusion

Genomic analysis of samples classified under the XBC and XBC.1 lineages collected in the Philippines from August to September 2022 suggests that these SARS-CoV-2 lineages are likely to be rooted in the country. In particular, the majority of these cases were detected in the adjacent southern Philippine regions of Davao (XBC) and Soccsksargen (XBC.1). The recombinant nature of these lineages was also observed, with regions corresponding to the Omicron BA.2 and Delta B.1.617.2 (Clade I) present in the sequences. This observation suggests that although the Omicron variant has been the predominant VOC in the Philippines since the beginning of 2022, lineages of the Delta variant are still circulating in the country albeit at much lower frequencies – making it possible for such recombinants to form. Evidence for this Delta-Omicron co-infection was also presented, further supporting the continuing co-circulation of these VOCs in the country and providing additional insights to the possible origins of the XBC recombinant lineage.

## Methodology

### SARS-CoV-2 sequencing, consensus sequence generation, and viral lineage designation

The reference-guided assembly, lineage assignment, and variant calling was done in the same method previously described in Tablizo et al [12]. For this analysis, the following tool versions were utilized: minimap2 v2.17 [13], Samtools v1.10 [14], iVar v1.3 [15], Pangolin v4.1.3 [16] [4], and MUMmer v4.0 [17] as implemented in the annotation tool RATT [18].

### Data visualization

Data visualization for daily confirmed COVID-19 cases (Fig 1) was generated using R 4.2.2 [19]with the packages ggplot2 3.3.6 [20] and ggpubr 0.4.0 [21]. The linelist of recorded SARS-CoV-2 cases from the Department of Health’s COVID Data Drop [22] was aggregated into daily cases and visualized with a histogram, then smoothed with a 7-day moving average.

The weekly proportions of detected lineages (Fig 3) were also visualized using ggplot2 3.3.6 after being filtered with tidyverse [23]. Samples that were not classified to a lineage were excluded from the visualization. In order to create a cleaner visualization, non-major Omicron sublineages (i.e. those that were not BA.2.3, BA.5.2, BA.5.10, or BA.2.3.20) were shown as “Other Omicron” and samples that were classified as hybrids but were not XBC or XBC.1 were shown as “Other Hybrids.” The sample patients’ regions were based on their listed address or the region of the collecting institution if the address is unavailable.

### Phylodynamic analysis

Phylogenetic trees were generated using Augur v11.3.0 and visualized using Auspice v2.32.1, both from the Nextstrain project [24]. As part of the canonical augur pipeline, quality filtering was performed before aligning the sequences using MAFFT [25]. Initial phylogenetic tree construction was performed using IQTREE [26] and was refined by rooting the tree to the oldest sample and by estimating confidence for inferred dates via marginal reconstruction using TreeTime v0.8.1 [27], dropping some samples as a part of the refinement process. TreeTime was also used to reconstruct ancestral traits and infer ancestral sequences as part of the augur pipeline employed for this research. All phylodynamic analysis is based on metadata associated with a total of 196 sequences available on GISAID from 2022-03-20 to 2022-10-18 and accessible at https://epicov.org/epi3/epi_set/230412yv?main=true.

### Mutation profiling

The summary plot for consensus mutations was produced from the iVar v1.3 [15] output using custom scripts written in Python. The key mutation profiles used for BA.2 and 21I Delta were obtained from outbreak.info [28] and covariants.org [29], respectively.

### Recombination analysis

Detection of recombination signals among the sequences was done separately using 3SEQ v1.7 [30] and Sc2rf [31]. 3SEQ run included specifying putative parent sequences from the local samples assigned to the Delta B.1.167.2 and Omicron BA.2 lineages, and the children assigned to the recombinant XBC and its sublineages. Visualization and the detection of the breakpoint regions is also observed using Sc2rf compared with the consensus sequences representing all other assigned XBC and XBC.1 local samples.

The plot for alternative allele fraction analysis was produced using custom scripts written in Python following the recombinant analysis visualizations in Bolze et al. [31]. Alternative allele frequencies for specific signature BA.2 and B.1.617.2 mutations were taken from the output of the ivar variants command in iVar v1.3 [15]. See the Data Availability section for the corresponding intrahost mutation files.

## Acknowledgments

We acknowledge the contributions of the various laboratories who have sent samples for whole genome sequencing and the Department of Health especially the Epidemiology Bureau for the national genomics biosurveillance effort and for making the sequence data publicly available. We also acknowledge the members of the Inter-Agency Task Force on Emerging Infectious Diseases (IATF) Task Force on COVID-19 Variants, Kenneth M. Kim, and Marc Jerrone R. Castro, whose foundational work in the Philippine SARS-CoV-2 genomic biosurveillance effort this paper is built upon. We gratefully acknowledge all data contributors, i.e., the Authors and their Originating laboratories responsible for obtaining the specimens, and their Submitting laboratories for generating the genetic sequence and metadata and sharing via the GISAID Initiative, on which this research is based.

## Data Availability

All genome sequences of the Philippine SARS-CoV-2 samples reported here are uploaded and shared publicly at the EpiCoV database of the Global Initiative for Sharing All Influenza Data (GISAID) database. For these sequences, and the sequences of the other international samples used in this analysis, their corresponding EPI ISL ID numbers are in the metadata file included in the Supplementary Data Table 2.

