## Supplementary Figure 1 for "Analysis of SARS-CoV-2 Recombinant Lineages XBC and XBC.1 in the Philippines and Evidence for Delta-Omicron Co-infection as a Potential Origin"

### SUPPLEMENTARY INFORMATION

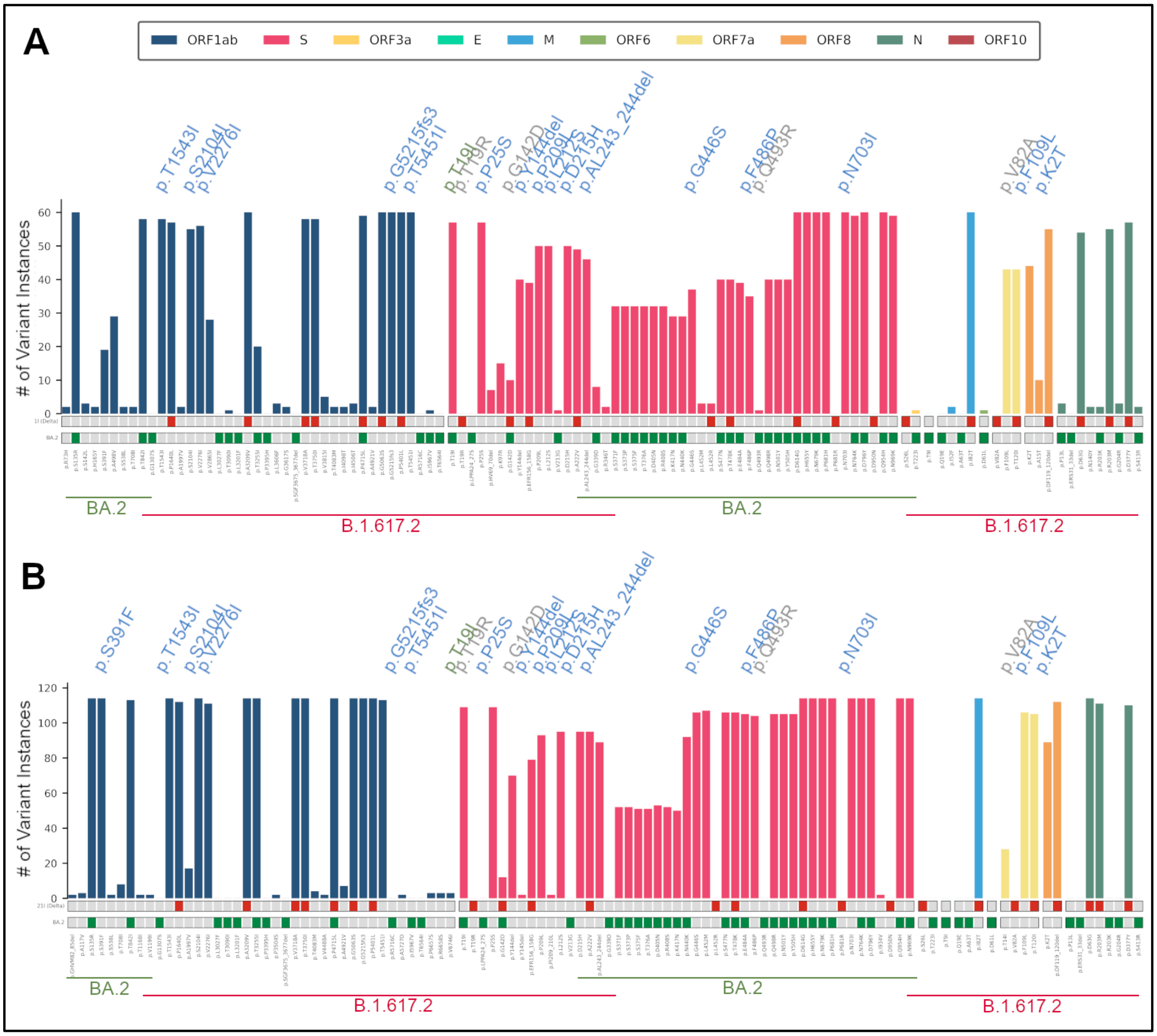


**Figure S1.** Consensus mutation profile of **(A)** 60 local XBC and **(B)** 114 local XBC.1 cases compared to the key mutations of BA.2 Omicron (green) and B.1.617.2 21I Delta Clade (red). These indicate the presence of mutations consistent with distinct sections of BA.2 and B.1.617.2. Highlighted are 18 distinct amino acid replacements (blue) and absences of key mutations (gray) compared to the BA.2 and B.1.617.2 sections. Additionally, though flanked by B.1.617.2 key mutations, the T19I Omicron mutation is found instead of the T19R Delta mutation. Based on these two mutation profiles, XBC.1 is found to have two more mutations than XBC: ORF1ab:S391F and S:L452M.
