## Supplementary Table 1 for "Analysis of SARS-CoV-2 Recombinant Lineages XBC and XBC.1 in the Philippines and Evidence for Delta-Omicron Co-infection as a Potential Origin"

### SUPPLEMENTARY INFORMATION

**Table S1.** Regions in the Philippines where the XBC and XBC.1 SARS-CoV-2 recombinant lineages have been detected as of October 14, 2022. Lineage classifications are based on the dynamic nomenclature system (Rambaut *et al.,* 2020) implemented in the Phylogenetic Assignment of Named Global Outbreak Lineage (Pangolin) tool.

| **Date** | **Region*** | | | **Date of Collection (dd/mm/yy)** |
|---|---|---|---|---|
| | **XBC** | **XBC.1** | **Total** | |
| International Travelers (IT) | 0 | 0 | 0 | - |
| National Capital Region (NCR) | 0 | 1 | 1 | 08/09/22 |
| Region 1 (Ilocos) | 0 | 0 | 0 | - |
| Region 2 (Cagayan Valley) | 0 | 1 | 1 | 20/09/22 |
| Region 3 (Central Luzon) | 0 | 0 | 0 | - |
| Region 4A (Calabarzon) | 0 | 3 | 3 | 26/08/22 - 12/09/22 |
| Region 5 (Bicol) | 0 | 0 | 0 | - |
| Region 6 (Western Visayas) | 0 | 0 | 0 | - |
| Region 7 (Central Visayas) | 0 | 1 | 1 | 19/09/22 |
| Region 11 (Davao) | 33 | 31 | 64 | 08/09/22 - 26/09/22 |
| Region 12 (Soccsksargen) | 22 | 76 | 98 | 25/08/22 - 26/09/22 |
| Region 13 (Caraga) | 3 | 1 | 4 | 22/08/22 - 03/09/22 |
| Cordillera Administrative Region (CAR) | 1 | 0 | 1 | 09/09/22 |
| Bangsamoro Autonomous Region in Muslim Mindanao (BARMM) | 1 | 0 | 1 | 10/09/22 |
| **OVERALL** | **60** | **114** | **174** | **22/08/22 - 26/09/22** |

* For local cases, ‘Region’ is based on the patient's address whenever available. If not, the region of the collecting institution is used. Due to limited sampling of cases for genomic biosurveillance, the values shown might not reflect the actual Omicron variant distribution and spread in the country.
