## Supplementary Table 2 for "Analysis of SARS-CoV-2 Recombinant Lineages XBC and XBC.1 in the Philippines and Evidence for Delta-Omicron Co-infection as a Potential Origin"

### SUPPLEMENTARY INFORMATION

Supplemental Table 2. List of Acknowledgements of authors, originating and submitting laboratories of SARS-CoV-2 sequences on GISAID.

#### SUPPLEMENTAL TABLE

**Data Availability**

GISAID Identifier: EPI_SET_230412yv
doi: 10.55876/gis8.230412yv

All genome sequences and associated metadata in this dataset are published in GISAID's EpiCoV database. To view the contributors of each individual sequence with details such as accession number, Virus name, Collection date, Originating Lab and Submitting Lab and the list of Authors, visit 10.55876/gis8.230412yv

**Data Snapshot**

- EPI_SET_230412yv is composed of 196 individual genome sequences.
- The collection dates range from 2022-03-20 to 2022-10-18;
- Data were collected in 7 countries and territories;
- All sequences in this dataset are compared relative to hCoV-19/Wuhan/WIV04/2019 (WIV04), the official reference sequence employed by GISAID (EPI_ISL_402124). Learn more at https://gisaid.org/WIV04.
